# Attractor competition enriches cortical dynamics during awakening from anesthesia

**DOI:** 10.1101/517102

**Authors:** Núria Tort-Colet, Cristiano Capone, María V. Sanchez-Vives, Maurizio Mattia

**Author notes:** **For correspondence:** (NTC). Equally contributing co-senior authors (María V. Sanchez-Vives, Maurizio Mattia).

## Abstract

Cortical slow oscillations (≲ 1 Hz) are a hallmark of slow-wave sleep and deep anesthesia across animal species. They arise from spatiotemporal patterns of activity with low degree of complexity, eventually increasing as wakefulness is approached and cognitive functions emerge. The arousal process is then an open window on the widely unknown mechanisms underlying the emergence of the dynamical richness of awake cortical networks. Here, we investigated the changes in the network dynamics as anesthesia fades out and wakefulness is approached in layer 5 neuronal assemblies of the rat visual cortex. Far from being a continuum, this transition displays both gradual and abrupt activity changes. Starting from deep anesthesia, slow oscillations increase their frequency eventually expressing maximum regularity. This stage is followed by the abrupt onset of an infra-slow (~ 0.2 Hz) alternation between sleep-like oscillations and activated states. A population rate model reproduces this transition driven by an increased excitability that brings it to periodically cross a critical point. We conclude that dynamical richness emerges as a competition between two metastable attractor states whose existence is here experimentally confirmed.

## Introduction

While asleep or under anesthesia, the brain generates slow oscillations (SO), a pattern of activity that alternates between periods of activity or Up states, and periods of silence or Down states (***Steriade et al., 1993; Chauvette et al., 2010; Sanchez-Vives et al., 2017***). SO constitute a multiscale phenomenon in the brain, since they initiate locally at the single-column level and then propagate as traveling waves across the cortical surface (***Massimini et al., 2004; Mohajerani et al., 2010; Huang, 2010; Ruiz-Mejias et al., 2011; Stroh et al., 2013***), recruiting the cortico-thalamo-cortical loop and other subcortical brain structures (***Steriade et al., 1993; Wolansky et al., 2006; David et al., 2013; Sheroziya and Timofeev, 2014; Amigó et al., 2015***). Such a global and stereotyped activity emerges from the synchronized cooperation between cortical assemblies oscillating between active (Up) and almost quiescent (Down) states (***Bazhenov et al., 2002; Compte et al., 2003; Destexhe, 2009***). Under these conditions brain activity displays a relatively low degree of complexity in dynamical terms (***Casali et al., 2013; Casarotto et al., 2016***). From a dynamical point of view, the local origin of slow oscillatory behavior has been hypothesized to result in assemblies of local neurons from the interplay between recurrent synaptic excitation and an activity-dependent adaptation or self-inhibition, giving rise to nonlinear dynamics that lead such networks to behave like relaxation oscillators (***Compte et al., 2003; Gigante et al., 2007; Curto et al., 2009; Mattia and Sanchez-Vives, 2012***).

The transition from the sleep to the awake state is known to bring the brain into a dynamical regime displaying an enriched repertoire of activity, thus leading to a high degree of complexity often associated with irregular firing “desynchronized” in time and space (***van Vreeswijk and Sompolinsky, 1996; Destexhe, 2009; Renart et al., 2010***). This dynamical richness allows the brain to perform state-dependent computations while maintaining the sensitivity to both sensory inputs and to a wide repertoire of inner mental states (***Buonomano and Maass, 2009; Deco et al., 2013***). Such complexity can emerge as an increase in the long-range functional connectivity of the cortical network (***Alkire et al., 2008; Bettinardi et al., 2015***) leading to restless collective dynamics, whenever a critical point of the phase diagram is approached (***Beggs, 2008; Deco et al., 2013; Tognoli and Kelso, 2014***). Changes in the local nonlinear dynamics of neuronal assemblies are also expected to play an important role in this brain state transition (***Bazhenov et al., 2002; Hill and Tononi, 2005; Curto et al., 2009***). Indeed, the onset of input-output nonlinear responses can be naturally expressed by the same local assemblies oscillating during slow-wave activity (***Curto et al., 2009; Mattia and Sanchez-Vives, 2012; Jercog et al., 2017***), giving raise to “flip-flop” units displaying a state-dependent response to fluctuating synaptic inputs (***McCormick, 2005; Linaro et al., 2011; Mattia and Sanchez-Vives, 2019***). Brain networks composed of such units can implement a wide range of computational capabilities ranging from enhancing or blocking sensorimotor processing, to integrating incoming information through long timescales (***McCormick et al., 2015; Huang and Doiron, 2017***). However, although the enhancement of the input-output nonlinear response is expected at the local assembly level (***Mattia et al., 2013; Reig et al., 2015; Zerlaut and Destexhe, 2017***), alternative interpretations of the experimental evidence about the synchronized-desynchronized state transition have been proposed. Indeed, the oscillatory behavior of local assemblies can also fade out due to a lowering of the excitability of the local networks, leading to a linearization of the input-output response (***Curto et al., 2009***). Due to these kind of ambiguities, the mechanistic root of the transition from sleep or deep anesthesia to wakefulness is still far from being fully understood.

Here, we focus on the dynamics underpinning the synchronized-desynchronized state transition at the local level of layer 5 (L5) neuronal assemblies, aiming at clarifying the scenarios mentioned before of an increase or a decrease of the network excitability. To this purpose, we recorded the neuronal activity in L5 from the primary visual cortex (V1) of rats during the transition from deep to light anesthesia; that is, from the slow oscillatory activity towards a pattern of activity characterized by the periodic appearance of micro-arousal periods (see below). The changes in the observed neuronal activity are fully captured by a population rate model in which SO spontaneously emerge from the interplay between recurrent synaptic excitation and activity-dependent self-inhibition (or adaptation) (***Compte et al., 2003; Mattia and Sanchez-Vives, 2012***). In this dynamical framework, the transition from deep to light anesthesia results from an increase in local excitability due to a decrease of the strength of adaptation and an increase of the external input, supporting the hypothesis of an input-output response amplification of L5 assemblies during the awakening.

## Results

In order to study the transition across different levels of vigilance in the cortex, five rats were deeply anesthetized and the local field potential (LFP) of a neural assembly of L5 in V1 was continuously recorded while the level of anesthesia progressively decreased (see Figure 1A and Methods). During the recovery from deep anesthesia, cortical rhythms underwent some gradual changes and eventually some abrupt ones, and this progressive transition was observed as different oscillatory patterns at the level of the LFP signal (Figure 1B).

**Figure 1.**
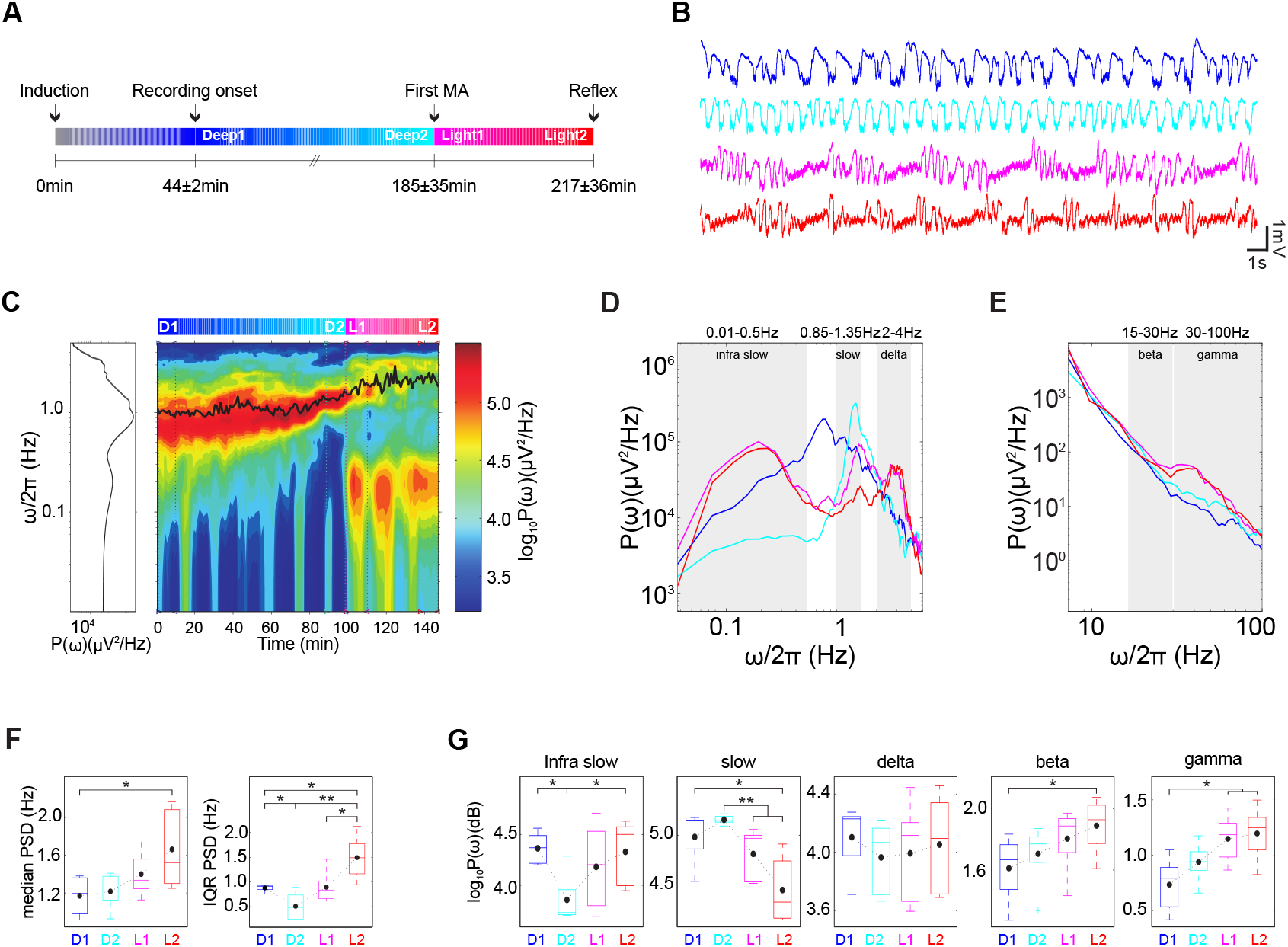
Rhythms involved in the transition from deep anesthesia towards wakefulness. **A**. Sketch of the time events used to define the different stages (D1, D2, L1 and L2) of the transition from deep anesthesia towards wakefulness (see Methods). Occurrence time of each event is averaged across animals (*n* = 5 rats) in the lower row. **B**. LFP traces extracted from the different stages of the transition in a representative recording (color coded as in the sketch in A). **C**. Spectrogram (0.01 – 5 Hz) of the LFP recorded during the experiment. Black superimposed trace corresponds to the median of the distribution of the PSD values at each time of the experiment. Blue, cyan, magenta and red vertical dashed lines indicate the boundaries taken in this example for each transition stage. Left panel shows the histogram of the spectrogram values throughout the experiment. **D**. and **E**. PSD (0.01 – 5 Hz and 5 – 200 Hz respectively) of the LFP recorded at different stages of the transition towards wakefulness (color coded as in the sketch in (A). Gray shaded areas indicate the frequency ranges used in G to compare the power at each frequency band in the different intervals. **F**. Population averages of the median PSD (left) and the IQR of the PSD (right) during each stage of the transition (color coded as in the sketch in A). **G**. Population averages of the power of different frequency bands during each stage of the transition (color coded as in the sketch in A). For panels F and G, black circles represent the mean of the distributions at each interval. On each box, the central line is the median (50*th* percentile), the edges of the box are the 25*th* and 75*th* percentiles and the whiskers extend to the most extreme data points not considered outliers. \**p* < 0.05, \*\**p* < 0.01, two-sample *t*-test, *n* = 5. MA: micro-arousal. D1: Deep1, D2: Deep2, L1: Light1, L2: Light2, LFP: local field potential, PSD: power spectrum density, IQR: Inter-quartile range.

As expected, at deep levels of anesthesia, L5 neurons showed SO at a frequency of 1.17 ± 0.21 Hz (mean ± standard deviation (SD), *n* = 5 rats) (***Steriade et al., 1993; Chauvette et al., 2011; Ruiz-Mejias et al., 2011***). The SO gradually became faster and more regular as the level of anesthesia decreased (Figure 1B) (see below). However, while reaching lighter levels of anesthesia, there was a characteristic abrupt onset of a new oscillatory pattern, in which the cortical SO was periodically interrupted by periods of low LFP amplitude: the micro-arousal states (***Bergmann et al., 1987; Watson et al., 2016***). The duration of these micro-arousal states ranged from a minimum duration of 0.68 ± 0.41 s to a maximum duration of 10.67 ± 5.87 s (mean ± SD across *n* = 5 rats). We describe this transition in detail in the following sections.

### Gradual and abrupt changes in the oscillatory patterns when anesthesia fades

In order to quantify the changes occurring during the recovery from anesthesia, we computed the power spectral density (PSD) of the LFP in 1-min windows throughout the time course of the experiment. As shown in the spectrogram of a representative recording in Figure 1C, during deep stages of anesthesia the dominant frequency was around 1 Hz (red and yellow areas). Then, as the anesthesia level decreased, the frequency of the SO gradually increased, as shown by the monotonic increase of the median PSD (Figure 1C). To perform population analyses across the subjects, we singled out four intervals of 10 min corresponding to different levels of anesthesia progressively advancing from deeper to lighter: Deep1 (D1), Deep2 (D2), Light1 (L1) and Light2 (L2) (see the anesthesia protocol in Figure 1A and Methods). During the deeper phases of anesthesia (D1 and D2), the median PSD frequency increased from 1.17 ± 0.21 Hz to 1.21 ±0.19 Hz (mean ± SD); and continued increasing during the lightest phases (L1 and L2) from 1.40 ± 0.24 Hz to 1.66 ± 0.42 Hz (Figure 1F, left panel).

This increase in the frequency of the SO was also observed as a shift in the peak at slow frequencies of the PSD of the D1 and D2 intervals (Figure 1D). At the same time, the spectrogram showed that the range of frequencies at which the recorded local network oscillated shrank as the level of anesthesia decreased (the red and yellow areas around 1 Hz in Figure 1C became concentrated in a smaller frequency range from D1 to D2). We quantified the reduction of the range of frequencies visited by the cortical network using the interquartile range (a measure of the variability of a distribution) which is the difference between the frequencies at the 75*th* and 25*th* percentiles of the PSD distribution computed at each minute during the course of the experiment. We found that, at population level, the range of frequencies started at 0.88 ± 0.07 Hz in D1 and dropped to a minimum in D2 with a range of 0.51 ± 0.28 Hz on average (Figure 1F, right).

These results suggest that during the first part of the transition from deep to light levels of anesthesia (from D1 to D2) the SO not only became faster but also more regular, since the interquartile range, representing the range of the frequencies visited around this median value, was reduced. This increased regularity of SO is highly suggestive that there is an optimal level of the network excitability to express this rhythm, as it has been reported for the cortex *in vitro* (***Sancristóbal et al., 2016***). We explore this quantitatively in our computer model (see below). When continuing towards lighter anesthesia (L1 to L2), the ongoing increase in regularity was disrupted by the sudden appearance of micro-arousals, which were identified as periods of low LFP amplitude that interrupted the regular SO every 5 s (0.2 Hz).

The presence of this new pattern of activity was associated with a peak in the low frequency range of the spectrogram (Figure 1C) and the PSD (Figure 1D), since the micro-arousal states appeared periodically at a frequency of around 0.2 Hz. The fact that another frequency in the lower region of the spectrogram became relevant would explain the larger interquartile range observed at the population level during the last phase of the transition: interquartile range increased up to 0.89 ± 0.33 Hz in L1 and finally to 1.49 ± 0.45 Hz in L2. (Figure 1F). We also observed that the power of the infra-slow oscillations (0.01 – 0.5 Hz) increased monotonically since the first appearance of a micro-arousal until the end of the experiment (*p* = 0.045, *two-sample t-test*, Figure 1G).

In order to investigate the nature of these periods of sustained asynchronous activity, we inspected the high-frequency content of the LFP signal recorded during different levels of (decreasing) anesthesia (from D1 to L2, see Methods). We found that the power of beta and gamma bands (intervals 15 – 30 Hz and 30 – 100 Hz, respectively) progressively increased during the transition from deep to light anesthesia, in particular during L1 and L2 (Figure 1EG Beta power in L2 was significantly higher than in D1 (*p* = 0.045, *two-sample t-test*). Gamma power in L1 had already increased significantly from D1 values and continued increasing during L2 (*p* = 0.033 and *p* = 0.027, respectively, *two-sample t-test*).

### Appearance of a new activated state

To describe the temporal patterns observed in the LFP during the fading of anesthesia, we extracted the multi-unit activity (MUA) signal from the raw signal, and singled out Up and Down states by thresholding the log(MUA) values (***Ruiz-Mejias et al., 2011***) (see Methods and Figure 2A). We detected the Down-to-Up state transitions occurring during 10-min intervals corresponding to different levels of decreasing anesthesia (D1, D2, L1 and L2, respectively). We then produced a raster of the MUA values during the Down-to-Up state transitions detected at each interval (Figure 2B).

**Figure 2.**
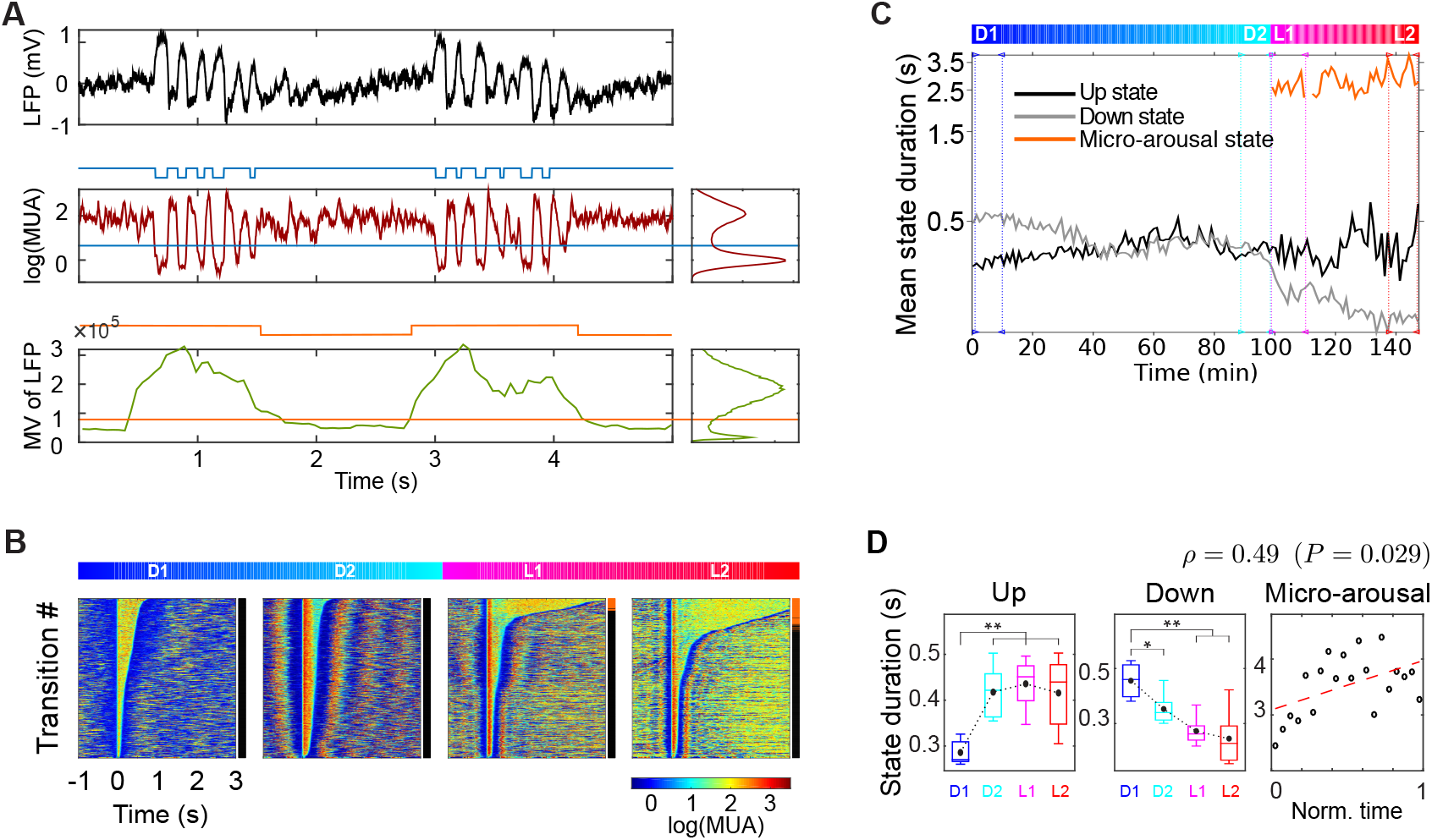
Changes in Up, Down and MA state duration across the experiment. **A**. LFP (top), MUA (middle) and moving variance (MV) of the LFP (bottom) versus time during a period in which slow oscillations alternate with micro-arousals. The panels on the right show the bimodal distribution of the histogram of the MUA or MV of the LFP values during all the experiment. Blue or orange horizontal lines show the thresholds used for Up/Down state or slow oscillations/micro-arousal detection in the case of MUA (blue) or LFP MV (orange) respectively. **B**. Raster plots of the MUA values during the Down-to-Up/micro-arousal state transitions, aligned at the Up/micro-arousal state onset and sorted by Up/micro-arousal state duration, in each stage of the transition from deep anesthesia towards wakefulness. Whether the transition is from a Down state towards an Up state or a micro-arousal is indicated at the right side of each raster plot with a black or orange stripe, respectively. **C**. Up, Down and micro-arousal state durations computed in 1-min time windows across the experiment. **D, left panels**. Population averages of Up and Down state durations in the different stages of the transition towards wakefulness (color coded as in the sketch in Figure 1A). **D, right panel**. Correlation between the duration of the micro-arousal state and its onset time (in the normalized time from L1 to L2). Boxplots as in Figure 1F,G. MUA: Multi-unit activity

Sorting the Up state onset times (i.e., the Down-to-Up state transitions) by Up state duration, we observed some interesting features concerning a new type of activity pattern. In particular, during the lightest phases of anesthesia (L1 and L2) a new kind of activated state with longer duration than Up states appeared: the micro-arousal state. To differentiate Up states from micro-arousals, which shared higher levels of MUA than Down states, we used a threshold on the moving variance of the LFP signal, since the LFP during micro-arousals showed very low variance compared to periods of SO (see Figure 2A and Methods). We then computed the mean duration of Up, Down and micro-arousal states in 1-min windows across the experiment (see Methods). We observed that while the Up state duration remained almost constant with a small tendency to increase, the Down state duration decreased monotonically, at some point converging to duration values very similar to those of the Up state, resulting into an increase in frequency and regularity of the SO (Figure 2C). Remarkably, a similar stability of the Up state duration and the changes of the Down state duration has been also found *in vitro* when the network excitability is modulated both by extracellular potassium concentration (***Sancristóbal et al., 2016***) and by applying exogenous electric fields (***D’Andola et al., 2018***) suggesting that regular SO is an attractive dynamical regime for the isolated cortex (***Sanchez-Vives et al., 2017***).

The population level analysis showed that Down states decreased monotonically from deep to light anesthesia, thus increasing the SO frequency (Figure 2D, middle panel), Up states remained rather constant even when they elongated slightly during the initial part of the transition (Figure 2D, left panel), and micro-arousals progressively elongated, resulting in a positive correlation between micro-arousal duration and its onset time (Figure 2D, right panel). This evidence was compatible with the scenario in which the increasing amount of time spent by the L5 network in the micro-arousals led to a duration decrease of the periods spent displaying coherent SO with increasing frequency.

### Optimal anesthesia level regularizing SO

Reduction of the level of anesthesia from D1 to D2 led to a more regular pattern of SO. However, at lighter levels of anesthesia, the appearance of micro-arousals altered the oscillatory pattern and thus weakened its regularity. To further investigate these changes in the regularity of SO, we computed the auto-correlogram of the MUA signal in 1-min time windows across the experiment. We found that when the level of anesthesia decreased, the time lag at which we found peaks of correlation increased (Figure 3A), indicating that the firing of the local network became more coherent as the level of anesthesia decreased from D1 to D2. Simultaneously, a minimum in the coefficient of variation (*c_υ_*) of both Up and Down state durations was reached (Figure 3B).

**Figure 3.**
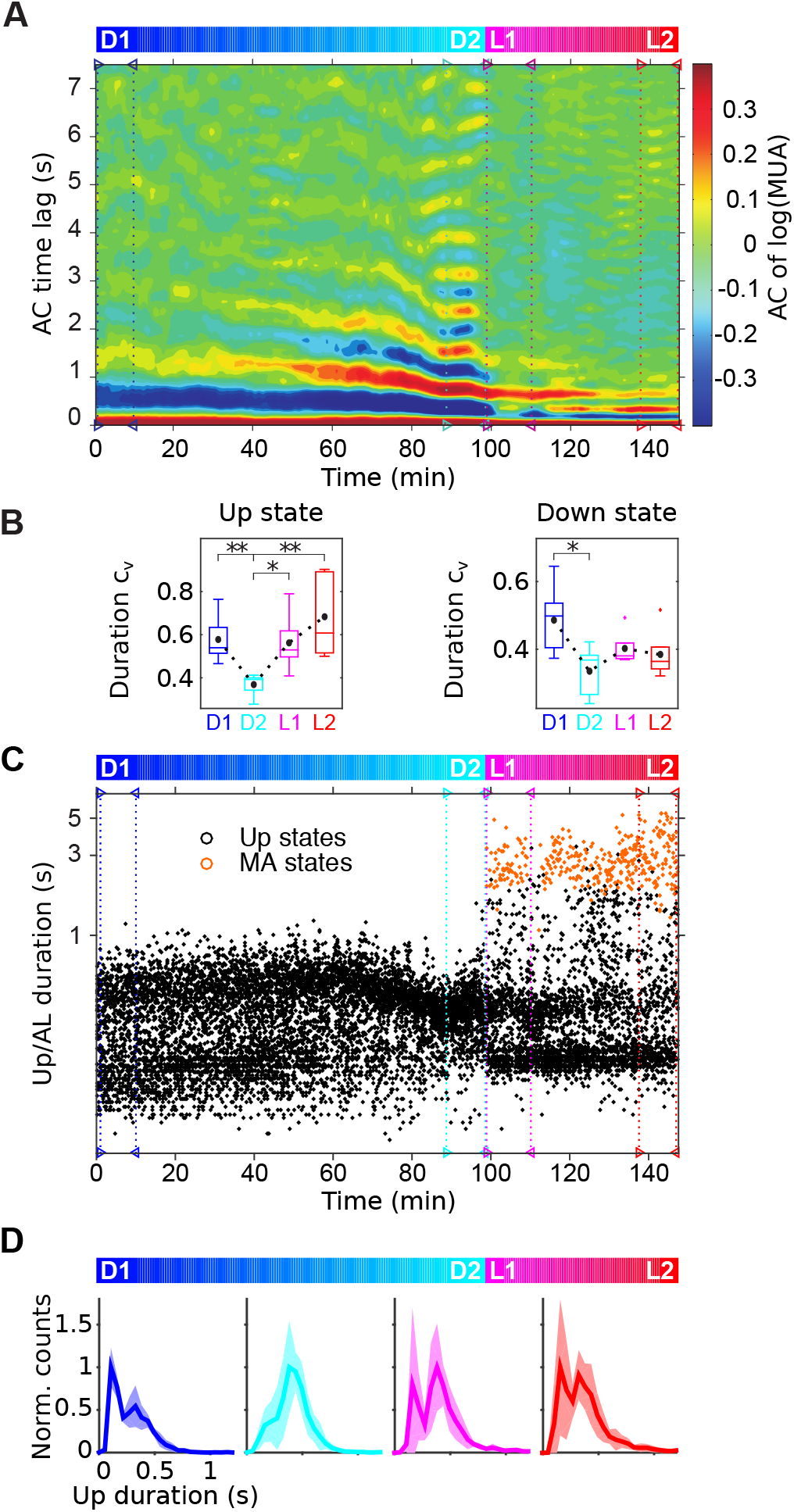
Optimal anesthesia level regularizing slow oscillations. **A**. Auto-correlogram of the log(MUA) signal computed at 1-min time windows across the experiment. Blue, cyan, magenta and red vertical dashed lines in A and C indicate the boundaries taken in this example for each transition stage. **B**. Mean coefficient of variation (*c_υ_*) of the duration of Up and Down states at each transition stage. **C**. Duration of Up states and micro-arousals (black and orange dots respectively) versus their onset time. **D**. Mean (solid line) and standard deviation (shadow) of the Up state duration distribution averaged across experiments in each stage of the transition towards wakefulness. Boxplots as in Figure1F,G. MA: micro-arousal.

We observed that Up state duration showed a bimodal distribution during deep anesthesia, with peaks at short (110 ms) and long (330 ms) durations (Figure 3D, D1). Interestingly, the fading of anesthesia occurring from D1 to D2 periods shrank the distribution of Up state durations. Indeed, right before the appearance of the first micro-arousal (Figure 3D, D2 period), the distribution of Up state durations became unimodal (with a preferred Up state duration slightly longer; 380 ms). On the contrary, when the micro-arousals started to appear (periods L1 to L2), the distribution of Up state durations became bimodal again (Figure 3C,D), breaking the MUA coherence (Figure 3A) and producing a decrease in the regularity of the Up state duration (*c_υ_*; Figure 3B).

Seen from the perspective of the dynamical system theory, such maximization of the regularity of SO, and thus the onset of more stereotyped Up states, is suggestive of an intriguing hypothesis: at this stage of the transition towards wakefulness (D2 period), our system (i.e., the probed L5 neuronal assembly) seems to pass through a region of its parameter space where only a dynamical state is available - a limit cycle - while at deeper and lighter anesthesia levels other attractor states are available to be visited. Under this hypothesis, the wandering across these attractors can account for the observed changes in the variability of Up and Down state duration.

### Micro-arousals are not longer Up states

It seems a logical question to wonder whether micro-arousals are just longer Up states. We investigated thoroughly this question, starting by evaluating the firing rate and the duration of both states during the final phases of the transition towards lighter anesthesia (L1 to L2). To determine the level of MUA of micro-arousals and compare it with that of the Up states, the asymptotic firing rate of Up and micro-arousal states was computed in 1-min time windows across the whole experiment (Figure 4A). While the asymptotic firing rate of Up states showed a tendency to increase as the anesthesia faded, that of the micro-arousals remained almost constant. In any case, those tendencies were not significant at the population level (Figure 4B). Interestingly, we observed in all cases that the mean asymptotic firing rate of micro-arousals occurring during the period from L1 to L2 was significantly lower than that of the Up states occurring during the same period (Figure 4C). This was already observed in the raster plots of the MUA around the transition towards the Up state or towards a micro-arousal state in the L1 and L2 intervals (Figure 2B).

**Figure 4.**
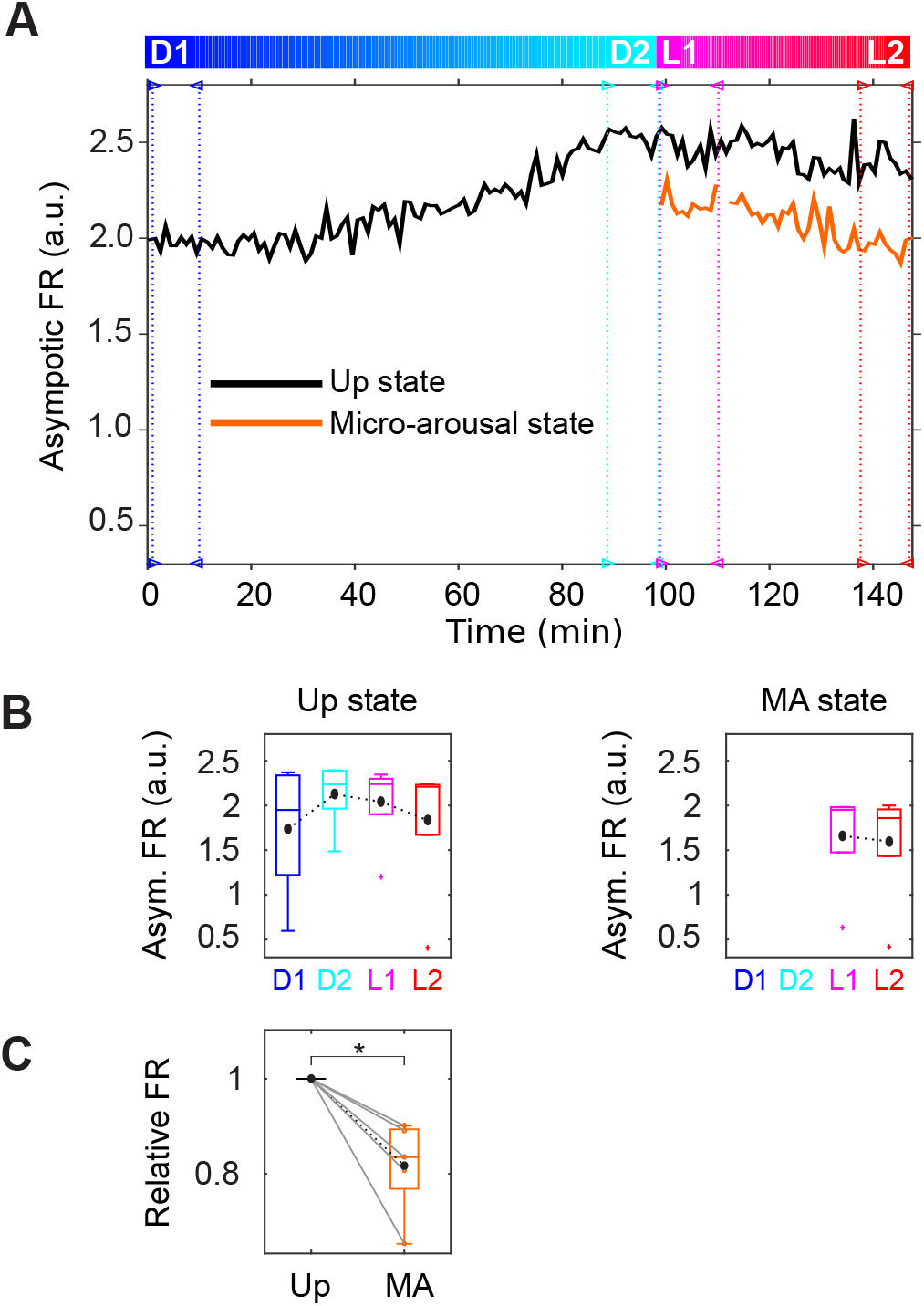
A micro-arousal is not merely a longer Up state: micro-arousals have lower firing rate compared to Up states. **A**. Up states and micro-arousals (black and orange respectively) asymptotic firing rate relative to the mean firing rate of the Down states computed in 1-min time windows across the experiment. **B**. Population average of the asymptotic firing rate during Up states and micro-arousals at each transition stage. **C**. Population average of the mean asymptotic firing rate of Up states (black) and micro-arousals (orange) during the part of the experiment in which slow oscillations and micro-arousals coexist (time from L1 to L2). For each subject, firing rate values were normalized by the value of asymptotic firing rate during Up states. Boxplots as in Figure 1F,G. MA: micro-arousal.

As both state duration and activity level of micro-arousals and Up states are different, one reasonable hypothesis is that their mechanistic, dynamical basis has different origins. In the following section we investigate this issue and use a model network of the L5 cell assembly to provide a coherent dynamical picture of the transition from deep to light anesthesia.

### State alternation onset in a model of the anesthesia fading out

The analysis performed on the data seemed to indicate that during the late phases of the arousal from deep anesthesia before awakening there was a competition between two types of dynamical regimes: the oscillatory (the slow alternation between Up and Down states) and the activated (micro-arousals).

In order to study in detail the mechanistic root of these dynamics, we developed a population rate model capable of reproducing the features of the lumped activity of the L5 assemblies we observed experimentally (Figure 5A). The aim of the model was to represent a network of a finite number of neurons displaying two metastable activity states at high (Up) and low (Down) firing rate *v*(*t*) due to a sufficiently strong recurrent synaptic excitation and a sigmoidal input-output gain function Φ(***I***), giving the asymptotic *v* of a neuron receiving a current ***I***. A quasi-regular alternation between Up and Down states was ensured by the presence of an activity-dependent adaptation level *a*(*t*) providing a proportional self-inhibitory current with strength *g_a_* (***Mattia and Sanchez-Vives, 2012***).

**Figure 5.**
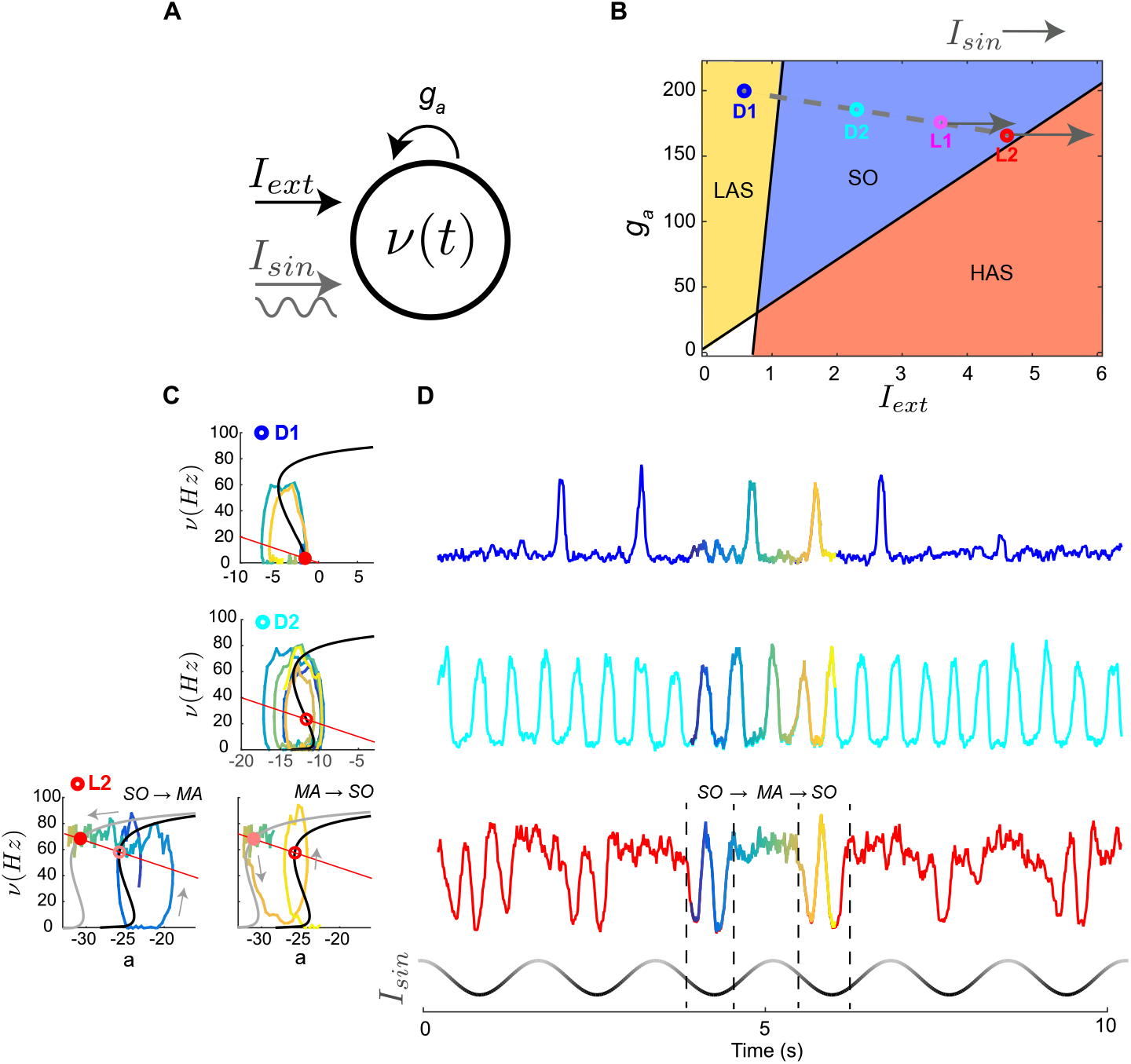
Modeling awakening from deep anesthesia in a neuronal assembly. **A**. Schematic diagram of the mean-field rate model of a cortical assembly reproducing *in vivo* recordings at varying levels of anesthesia. The model accounts for two variables: the firing rate *v*(*t*) and an activity-dependent self-inhibition (adaptation) *a*(*t*) with strength proportional to the coupling variable *g_a_*. The excitability of the system is modulated by an external input current which can be oscillatory (***I**_sin_*) or constant in time (***I**_ext_*). **B**. Bifurcation diagram (***I**_ext_, g_a_*) summarizing all the dynamical regimes the model is able to express: a stationary state at low (high) firing rate values in the LAS (HAS) region and an oscillatory state (stable limit cycle) in the SO region. In the white region both high and low stationary regimes coexist (fully bistable regime). The dashed gray line is the trajectory modeling the fading of anesthesia, with different stages (D1,D2,L1 and L2) depicted as colored circles. Horizontal arrows represent onset and amplitude of the oscillatory external input current, ***I**_sin_*. **C**. Phase portraits (*a, v*) of the modeled network representing different anesthesia levels: D1, D2 and L2 from top to bottom, respectively. Black and red solid lines, nullclines of *v* (*dv/dt* = 0) and *a* (*da/dt* = 0), respectively, which intersections represent fixed-points of firing rate and adaptation level (red circles). Example trajectories from the periods of time highlighted in (D) are plotted in the same color code as in D. For L2 two samples trajectory are reported, to stress out the two transitions: the transition from SO to a micro-arousal and the transition from a micro-arousal back to SO. For the lightest level of anesthesia, two phase portraits indicate the different positions of the sigmoidal nullcline of *v* produced by different values of the external oscillatory input ***I**_sin_* that lead to SO (top) or micro-arousal (bottom) periods. **D**. Representative simulations of the network model activity *v*(*t*) of the three dynamical regimes highlighted in (C). Periods of time used in (C) are those plotted here with varying hues. Last row depicts the time course of the oscillatory input, ***I**_sin_* that produced the transitions from the limit cycle (SO) to the fixed point (micro-arousal) in the lighter level of anesthesia. SO: slow oscillations, MA: micro-arousal, LAS and HAS: low and high attractor state, respectively.

In addition, the activity *v*(*t*) of the network was modulated by an external input current ***I**_ext_* representing the net excitatory input due to the presynaptic activity coming from neurons external from the modeled network, and additionally by an oscillatory (sinusoidal) external input (***I**_sin_*) that under suitable dynamical conditions could be turned on (see Methods) to reproduce the SO-micro-arousal alternation observed under light anesthesia (L1-L2). To include the endogenous fluctuations of v(t) due to the unavoidable statistical variability of the spikes emitted per unit time in a network composed of a finite number of neurons (***Mattia and Del Giudice, 2002***), we also added a white noise in the input current ***I*** with appropriate intensity (see Methods).

The state of this two-dimensional system is a point in the phase plane (*a, v*) which moves in time forming specific trajectories. Due to the nonlinearity in amplifying input current ***I*** and the inhibitory feedback *g_a_ a* underlying spike-frequency adaptation, this kind of system can display a rich repertoire of dynamical regimes (***Strogatz, 2014***). Indeed, by inspecting the bifurcation plane defined by the adaptation strength *g_a_* and the excitability level associated with the current ***I**_ext_*, regions with both attracting fixed (equilibrium) points and with periodic orbits (limit cycles) can be identified (Figure 5B) (***Mattia and Sanchez-Vives, 2012***). More specifically, when the inhibitory *g_a_* or the excitatory ***I**_ext_* component is predominant, a stable fixed point (attractor) arises at relatively low (LAS, or low attractor state, top-left corner) or high (HAS, high attractor state, bottom-right corner) firing rate, respectively. When self-inhibition is sufficiently balanced by the network excitability, an unstable fixed point emerges giving rise to an oscillatory regime (SO region), related to a limit cycle.

To mimic the fading out of anesthesia, we implemented a rather specific increase of the excitability degree in the model by simultaneously increasing the external current ***I**_ext_* and reducing the strength of adapatation *g_a_*. We took into account that ketamine is an NMDA receptor antagonist and as such reduces the glutamatergic synaptic input from upstream cortical and subcortical areas (***Alkire et al., 2008; Brown et al., 2011***). This, together with a reduced excitatory noradrenergic input to infragranular layers expected when the locus coeruleus (***McCormick, 1992***) is inhibited by medetomidine (***Brown et al., 2011***), led us to impose a low initial value for ***I**_ext_*, eventually increasing it as the anesthesia fades out. As in the work of ***Hill and Tononi*** (***2005***) and ***Destexhe*** (***2009***), the reduction of the adaptation-related feedback *g_a_* was implemented to mimic the effect of lowered acetylcholine (ACh) levels induced by ketamine anesthesia (***Alkire et al., 2008***). Indeed, low levels of ACh lead to a larger slow *Ca*^2+^-activated ***K***^+^ current enhancing the spike-frequency adaptation of pyramidal neurons (***McCormick, 1992***). Such strong adaptation is reduced as the levels of ACh return to baseline, eventually contributing to the increase in cortical excitability as anesthesia fades. As a result, the the gradual changes of *g_a_* and ***I**_ext_* are shown in Figure 5B as a gray-dashed trajectory moving from the deepest anesthesia level D1 to the lightest L2 we singled out in the experiments. Moving the system along this trajectory, all the dynamical conditions observed in the different stages of anesthesia were reproduced: irregular SO in D1 (in LAS close to edge with SO, Figure 5B), regular SO in D2 (SO region, Figure 5B) and the SO-micro-arousal alternation in L1-L2 (periodically crossing the edge between SO and HAS, Figure 5B).

In order to visualize how the different dynamical regimes were generated by this simple bidimensional system, we plotted the nullclines of *v*(*t*) and *a*(*t*) (black and red curves in Figure 5C, left panels) for different levels of anesthesia (D1, D2 and L2). Nullclines indicate where the time derivatives of the variables vanish (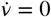 and *ȧ* = 0, respectively). The crossings of the nullclines, the fixed points, are the points in the parameter space where both derivatives are zero. The top panel of Figure 5C represents the nullclines of the system in D1 showing that the Down state is a stable fixed point and the irregular occurrence of Up states is due to the aforementioned endogenous fluctuations (Figure 5D, top panel). In D2 (Figure 5C, central panel) the fixed point was unstable, indicating the presence of a limit cycle that leads to regular oscillations (Figure 5D, central panel) as observed in the experimental data (Figure 1B, second row). When the oscillatory external input (***I**_sin_*) was turned on (during L1-L2), the position of the *v*-nullcline changed over time. An increase (decrease) in ***I**_sin_* shifted the nullcline towards the left (right) in the (*v, a*) plane, producing the gray (black) curves in Figure 5C (bottom panels). For minimal values of ***I**_sin_*, an unstable fixed point (limit cycle) produced a periodic orbit around the crossings of the *v*-nullcline (black) and the *a*-nullcline (red) giving rise to the oscillatory regime. For maximal values of ***I**_sin_*, a stable fixed point was created at the crossing between the two nullclines at a high value of *v*, leading to a HAS regime. Thus, during the latter phases of the transition from deep to light anesthesia, the presence of this oscillatory external input ***I**_sin_* induced a periodic subcritical Hopf transition between the limit cycle (SO region) and the high activity attractor (HAS region), resembling the SO-micro-arousal alternation observed in the experiments (Figure 5D, bottom panel).

The synthetic MUA obtained in the simulation displays a time course of its Fourier-frequency content which is remarkably similar to the one recorded during the experiments (compare the spectrograms of Figures 1C and 6A). Indeed, the modeled MUA reproduced the increase in the frequency of the SO and the appearance of a peak in power at ~ 0.2 Hz. Moreover, the evolution of the duration of the Down, Up and micro-arousal states during the transition from deep to light anesthesia (Figure 6B) was fully consistent with experimental observations (Figure 2C,D): mean Down state duration (gray curve) decreased, mean Up state duration (black curve) increased and mean micro-arousal duration (orange curve) increased from L1 to L2 periods. Finally, by comparing the MUA level of Up states and micro-arousals (during L1 and L2 light anesthesia periods), we found that the micro-arousals had, on average, lower activity levels than the Up states (Figure 6C) as seen in the experiments (Figure 4C).

**Figure 6.**
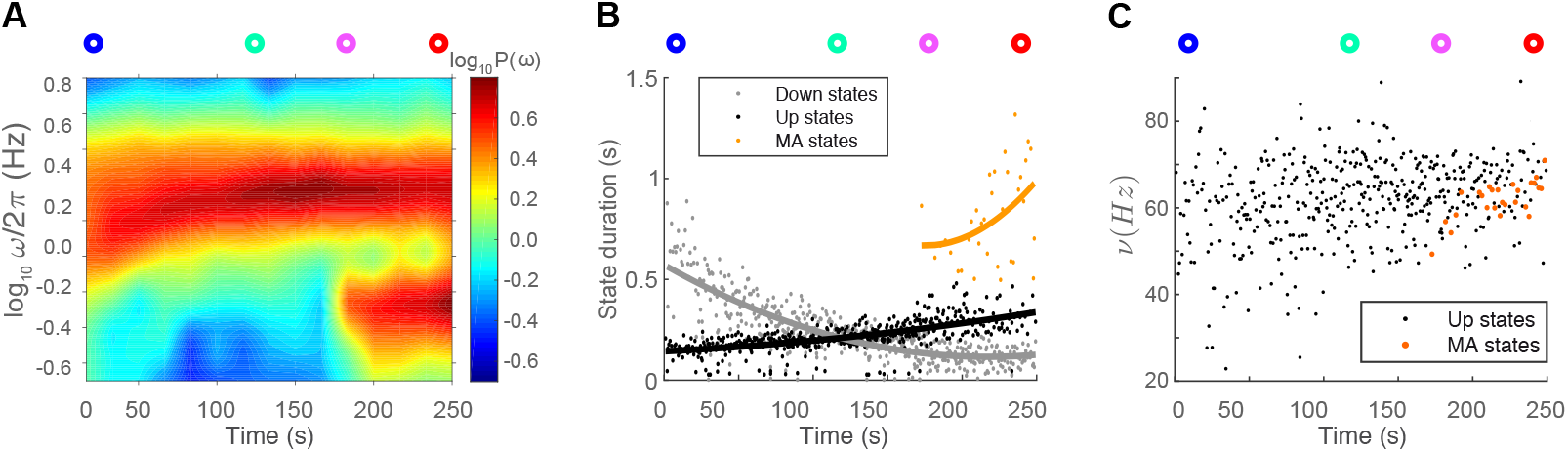
Modeling awakening from deep anesthesia in a neuronal assembly. **A**. Spectrogram of the firing rate *v*(*t*) as the network parameters are varied following the trajectory shown in (Figure5B). **B-C**. State durations (B) and average firing rate (C) of detected Up states (black), micro-arousals (orange) and Down states (gray) across a simulation in which the entire trajectory in the diagram (Figure5B) is crossed at constant speed. MA: micro-arousal.

### Footprints of the modulation of attractor stability in model

We further investigated the dynamics underlying the switch from the oscillatory to the stationary state, that we identified by the Down-to-micro-arousal transition. We first analyzed the time course of the network’s activity around the Down-to-Up and Down-to-micro-arousal transitions and found that the former had a larger MUA amplitude, involving a lower Down state MUA level and a higher maximum MUA during the Up state with respect to the latter (Figure 7A). Moreover, the transitions towards micro-arousals displayed a larger degree of variability when compared to the transitions towards Up states (shaded area in Figure 7A).

**Figure 7.**
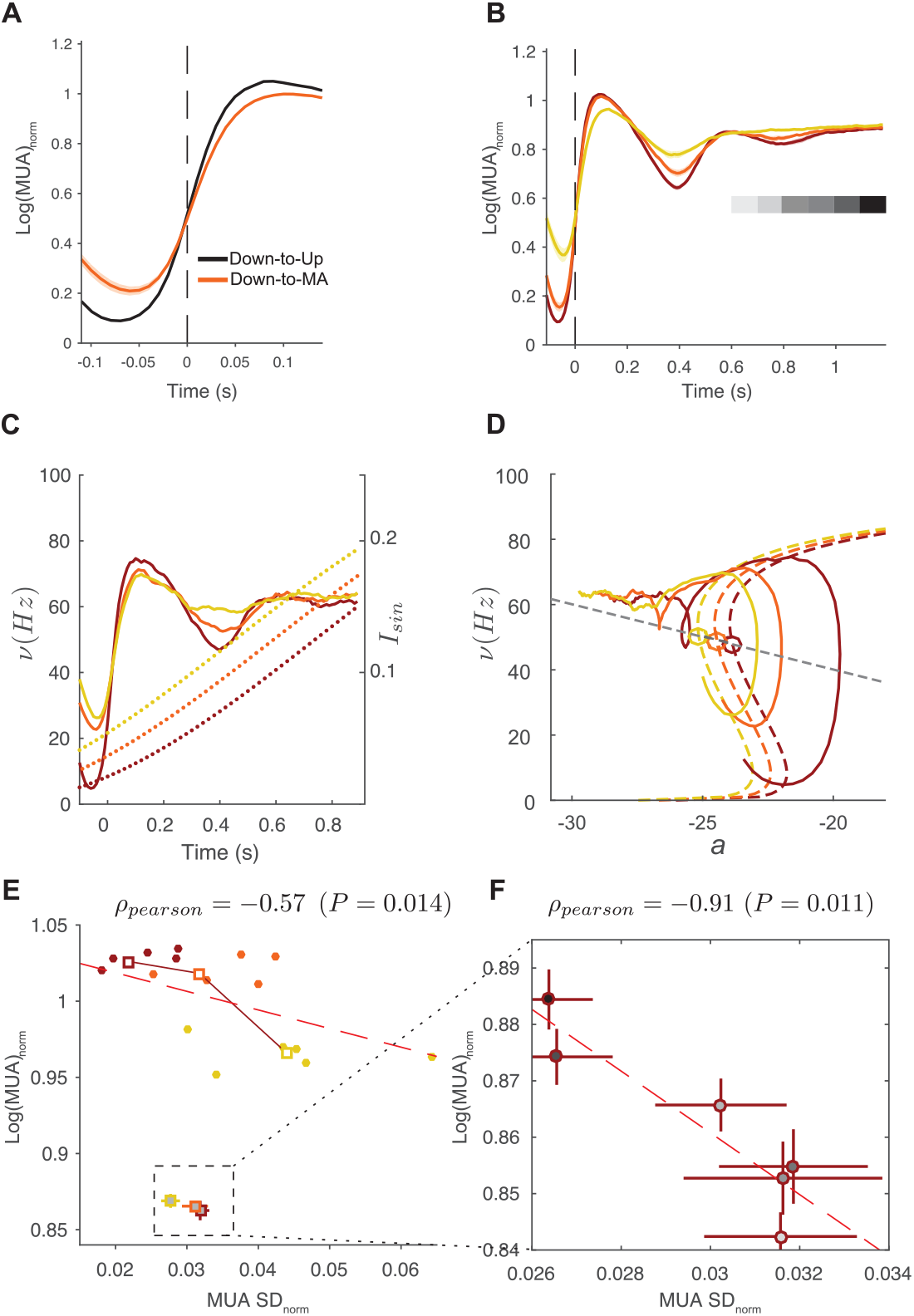
Expected fingerprints of attractor dynamics from the modeled neural assembly. **A**. Average firing rate *v* (i.e., modeled MUA) of detected Down-to-Up state and Down-to-micro-arousal transitions (black and orange, respectively) during the simulated L1 anesthesia level. Shadings, MUA standard deviation (SD). **B**. Average MUA of micro-arousals centered around the Down-to-micro-arousal transition onset pooled into three different groups with increasing firing rate at the end of the preceding Down state (from light to dark orange). Gray shaded boxes indicate six consecutive time intervals of 200 ms of increasing MUA following the micro-arousal onset. **C**. Average MUA of mircro-arousals centered around Down-to-micro-arousal transitions, pooled into three different groups with increasing firing rate at the end of the preceding Down state (solid lines from light to dark orange). Dotted lines in light to dark orange represent the time course of the oscillatory external input ***I**_sin_* averaged over micro-arousals in the respective pool. **D**. Nullclines of the two-dimensional system variables: *a* (gray dashed line) and *v* (light to dark orange dashed lines). The latter are referred to the minimum value of external drive received in that group of micro-arousals. Solid lines represent the trajectory that each group of micro-arousals would follow in the (*v, a*) plane during the transition represented in panel E. **E**. Pearson correlation between mean and SD of normalized log(MUA) computed at the peak of the waveform average for each group of micro-arousals in B. Squares and dots, average and single values from five different simulations, respectively. Red dashed line, linear fit of the data from all simulations (dots). Dashed box inset: mean and SD of MUA computed during the late part of the micro-arousal pointed out with the gray boxes in B. **F**. Pearson correlation between mean and SD of normalized log(MUA) computed during the six time intervals coded as different gray intensities during the waveform average for each group of micro-arousals in B. Error bars, SD of values estimated from five different simulations with different finite-size noise realizations. SD: standard deviation

To explore whether the variability among micro-arousals could be generated by differences in the pre-transition activity, Down-to-micro-arousal transitions were sorted by the level of activity in the previous Down state and grouped into three clusters of equal size (Figure 7B). We found that the cluster characterized by a lower level of activity during the previous Down state (dark orange) reached higher values of activity during the beginning of the micro-arousals.

In our model the presence of Down states with different activity levels is due to the slow oscillating input ***I***_*sin*_, and such activity levels are directly proportional to the value of ***I***_*sin*_ at the moment of the Down-to-micro-arousal transition (see Figure 7C, where the value of ***I***_*sin*_(*t*) for different clusters is reported in dashed lines).

The presence of different micro-arousals can be represented as different trajectories in the (***v, a***) plane (Figure 7D, same colors as in panels B,C). The dark orange trajectory, where the value of ***I***_*sin*_ is the smaller, is very close to the limit cycle describing the SO regime. The other trajectories (lighter orange) are less close to such a limit cycle, showing that higher values of ***I***_*sin*_ are related to more destabilized stages of this attractor.

In order to quantify the destabilization of the limit cycle, we evaluated the size of the fluctuations at the beginning of the micro-arousals by computing the inter-state standard deviation (SD, see Methods for details) for each group of micro-arousals. It is indeed known that more stable fixed points lead to smaller fluctuations in the dynamics, since their basin of attraction is narrower and hence the restoring force is stronger (***Mattia and Sanchez-Vives, 2012; Mattia et al., Under review***). We found an anti-correlation (***ρ***_*pearson*_ = −0.57,***P*** = 0.014) between the level of activity reached during the transition (initial peak) and the SD at the peak of the micro-arousals (Figure 7E).

As previously discussed, the transition to the micro-arousals is due to the creation of a stable fixed point (strong red circle in Figure 5C, bottom-left panel). In order to evaluate the stability of this new fixed point, we looked at the time course of the averaged profile of MUA during micro-arousals (Figure 7B). We observed that there was a strong effect of adaptation that reduced the firing after the initial peak of activity, similar to that seen during the offset of Up states (***Sanchez-Vives and McCormick, 2000; Compte et al., 2003***). However, the activity did not drop completely, but rather slowly increased again for a few hundred milliseconds (gray shaded squares in Figure 7B).

We evaluated the MUA level and the SD at consecutive time windows of 100 ms starting at 600 ms after the micro-arousal onset (gray shaded squares in Figure 7B) and found (Figure 7F) that they were strongly anti-correlated (***ρ***_*pearson*_ = −0.91, ***P*** = 0.011), indicating that while the level of activity increased, the size of fluctuations (SD) decreased, reflecting the presence of a fixed point whose stability was progressively increased.

### Evidence of attractor stability changes in experiments

Our model predicted the presence of more than one attractor whose stability would be modulated as anesthesia fades out by the oscillating external input, leading to the dynamics shown in Figure 5. Suggestions for the physiological basis of this external input are discussed below. In order to further strengthen the power of our model and its underlying assumptions, we tested these predictions using the experimental data. First, we compared the average profile of the normalized log(MUA) for the Down-to-Up state transition and the Down-to-micro-arousal state transition (black and orange curves in Figure 8A). The profiles were centered around the transition time and averaged over the experiments. Comparing this result with that in Figure 7A, it was apparent that the amplitude of the initial MUA peak was larger in the Down-to-Up state transition, as the model predicted.

**Figure 8.**
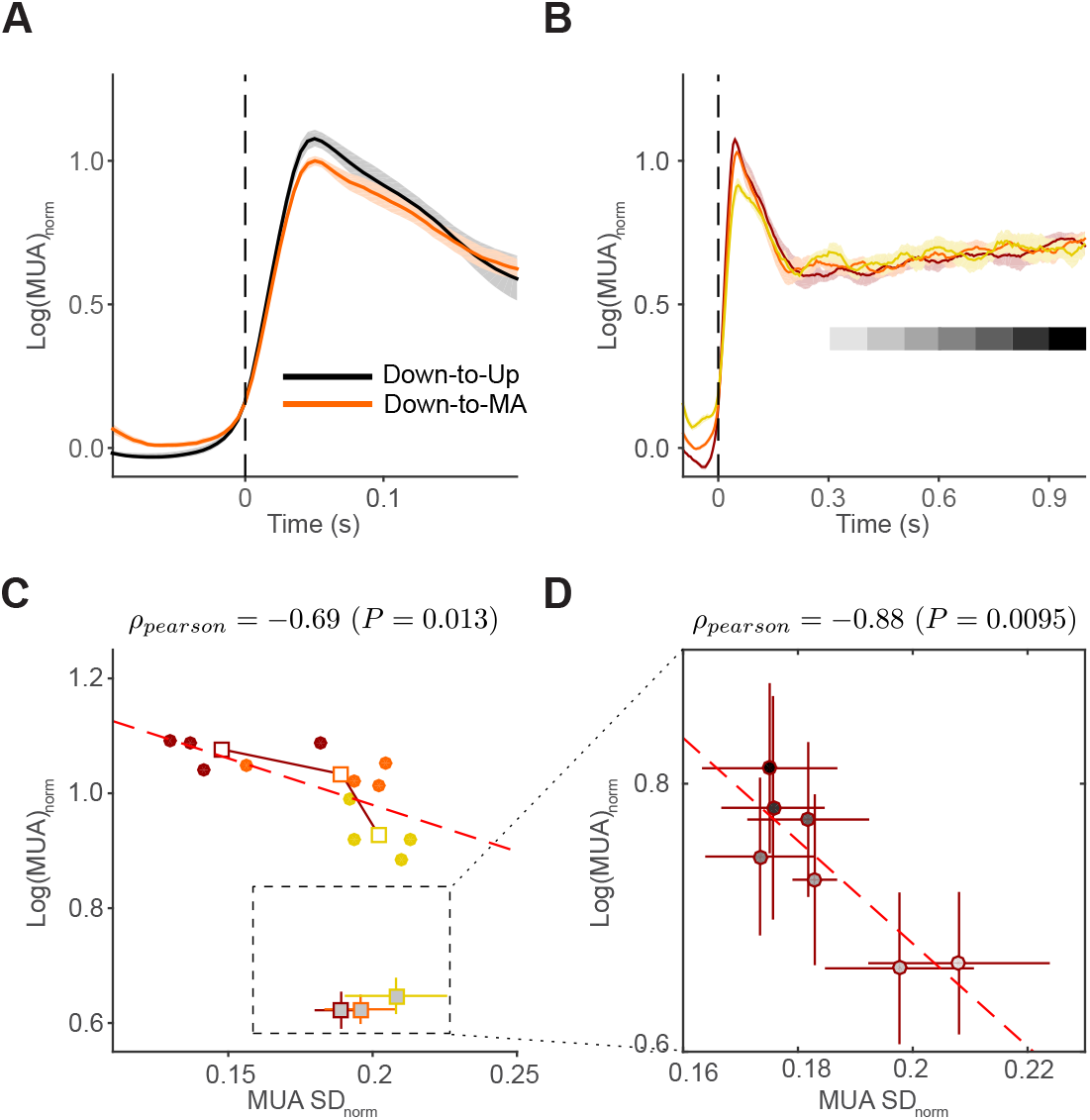
*In vivo* micro-arousals are attractors with exogenously modulated stability. **A.** Average log(MUA) during Down-to-Up and Down-to-micro-arousal transitions (black and orange, respectively) detected during the light anesthesia level (from L1 to L2). Shaded area: MUA standard deviation (SD). **B.** Average log(MUA) of micro-arousals centered around the Down-to-micro-arousal transition onset pooled into three different groups with increasing firing rate (from light to dark orange) at the end of the preceding Down state as in Figure 7B. Gray shaded boxes indicate seven consecutive time intervals of 100 ms of increasing MUA following micro-arousal onset. **C.** Pearson correlation between mean and SD of normalized log(MUA) computed at the peak of the waveform average for each group of micro-arousals in B. Squares and dots, average and single values from four different experiments, respectively. Red dashed line: linear fit of the data from all subjects (dots). Dashed box inset: mean and SD of log(MUA) computed during the late part of the micro-arousal pointed out with the gray boxes in B. **D.** Pearson correlation between mean and SD of normalized log(MUA) computed during the seven time intervals coded as different gray intensities during the waveform average for each group of micro-arousals in B. Error bars, SD of values estimated across four different experiments. MUA is normalized to the maximum activity measured during micro-arousals. SD: standard deviation, MA: micro-arousal.

In order to investigate the sources of variability among Down-to-micro-arousal transitions, micro-arousals were sorted by the level of activity in the previous Down state and grouped into three clusters of equal size, as we did with the simulated data. The within-cluster average profiles around the transition shown in Figure 8B confirmed that the cluster characterized by a lower MUA level during the Down state before the transition (darker orange) had a higher peak in MUA during the micro-arousal. As done for the simulated data, we evaluated the SD of the MUA at the peak of activity during the micro-arousal in each cluster and found a significant anti-correlation (***ρ***_*pearson*_ = −0.69, ***P*** = 0.013) between the mean MUA value and the SD of MUA across different clusters (Figure 8C), enforcing the hypothesis of a progressive destabilization of the SO. We observed in all clusters that the initial drop of activity after the peak was followed by a slow increase in the MUA level during the late part of the micro-arousal (Figure 8B). Confirming the model predictions, the increase in the MUA level was accompanied by a decrease in the SD, showing a significant anti-correlation (***ρ***_*pearson*_ = −0.88, ***P*** = 0.0095; Figure 8D) that confirmed the progressive stabilization of the fixed point for the micro-arousal predicted by the model.

Our results indicate that the dynamics of the cortical assemblies during the recovery from deep anesthesia are governed by the competition of two attractor states, a limit cycle that leads to oscillatory activity and a stable fixed point at high values of firing rate that leads to asynchronous micro-arousal periods, and that this competition is modulated by an external oscillatory input.

## Discussion

Our results indicate that far from being a continuum, the transition from deep anesthesia to wakefulness is characterized by both gradual and abrupt changes in the LFP and MUA dynamics. This can be fully explained as a progressive imbalance between two competing attractor states. Crucially, we show that the transition from the oscillatory (i.e., slowly alternating metastable Up and Down states) to the asynchronous regime does not emerge as a progressive morphing of one state (the SO) into another (the micro-arousal state), but rather passes through a slow, periodic (~ 0.2 Hz) bouncing between SO and micro-arousals in which the latter occupies a progressively increasing fraction of the cycle, eventually surviving as the only available network state when wakefulness is approached (Figure 2D). This scenario is qualitatively different from the one emerging in similar nonlinear systems aiming at modeling spontaneous brain activity which, in the absence of external inputs, restlessly move around a mean-field critical point, eventually giving rise to a scale-invariant distribution of the micro-arousal duration (***Deco et al., 2013; di Santo et al., 2018***). The SO-microarousal alternation is also not captured by cortical network models in which the population activity is represented by a single stochastic state-variable (***Deco et al., 2009***). Indeed, in these models, Up and Down states are metastable attractors and during light anesthesia the network hops between them randomly in time, which is incompatible with the coherence in time of the 0.2 Hz rhythm observed in our experiments (Figure 1C). Furthermore, in this framework Up states would resemble small fragments of wakefulness (***Destexhe et al., 2007***), as they represent a continuum with the activated micro-arousal state. In contrast, the results we report for anesthetized rats seem to challenge this view, since Up states and micro-arousals show markedly different features both in terms of MUA profiles in time (Figure 8A) and in the intensity of the activity (Figure 4).

Before crossing such a critical point, as the anesthesia fades out from the deepest level (D1), the SO become maximally frequent and regular as level D2 is approached (Figure 1C,F). This increase in frequency and regularity of the SO is consistent with previous findings indicating that, at the macroscale, coherence between distant cortical areas at the frequency of the SO increases as the level of anesthesia decreases and that the peak of correlation shifts to slightly higher frequencies (***Bettinardi et al., 2015***). Here, the increasing frequency of SO is due to the monotonic decrease of the Down state duration, supporting the hypothesis of an increased excitability of L5 assemblies which results in a destabilization of the low-firing state (Figure 2C,D). As the Down state becomes less stable, SO reach their maximum regularity (minimum *c_ν_* of both Up and Down state durations, D2 stage in Figure 3B) signalling the absence of other competing attractor states (low and high asynchronous states, Figure 5B). The strong stability of SO at this level of anesthesia has to be considered a possible pathological condition. Indeed, under natural sleep the variability of Up state duration is markedly larger (***Watson et al., 2016***), although under the deepest non-rapid eye movement (NREM) sleep stage regularity measured as the height of the 1 Hz peak in the EEG power spectrum is at a maximum ((***Amzica and Steriade, 1997; Borbély et al., 1981***). In addition, such enhanced stability implies a resilience to external perturbations (***Sanchez-Vives et al., 2017***), and as a result, a low perturbational complexity (***Casali et al., 2013***) should be measured.

The emergence of different dynamical regimes in the probed L5 assemblies was fully captured by a minimal population rate model. Under mean-field approximation, the model provides an effective description of a network of excitatory neurons with spike-frequency adaptation (***Gigante et al., 2007; Mattia and Sanchez-Vives, 2012***). To mimic the transition from irregular SO (D1) to the onset of micro-arousals (L1-L2), we implemented a rather specific increase of the excitability degree in the model by simultaneously increasing the external current ***I_ext_*** and reducing the adaptation strength ***g***_*a*_. In this way, we modeled the effects of ketamine (an NMDA receptor antagonist) and medetomidine (inhibiting the noradrenergic system) both affecting the excitatory input ***I***_*ext*_ from upstream cortical and subcortical areas (***Alkire et al., 2008; Brown et al., 2011***). On the other hand, as in ***Hill and Tononi (2005)*** and ***Destexhe (2009)***, the lowering of the adaptation-related feedback ***g***_*a*_ mimicked the effect of the reduction of the ACh levels induced by ketamine-anesthesia (***Alkire et al., 2008***) (see Results for further details). A qualitatively similar picture can be obtained by increasing the strength ***J*** of the recurrent synaptic excitation, which, in order to maintain the spontaneous alternation between Up and Down states, cannot be lowered below a certain value (***Compte et al., 2003; Gigante et al., 2007; Mattia and Sanchez-Vives, 2012***). Indeed, under stationary conditions, the input-output gain function Φ(***I***) is proportional to ***J***−***g***_*a*_***τ***_*a*_, such that reducing the spike-frequency adaptation strength ***g***_*a*_ is equivalent to increasing the synaptic self-excitation ***J***. This scenario differs from the one proposed by (***Bazhenov et al., 2002; Deco et al., 2014; Krishnan et al., 2016***), where, in addition to an increased excitability of the cortical network due to decreased-leak ***K***^+^ currents (equivalent to an increase of ***I***_*ext*_), a reduction of the pyramidal-to-pyramidal glutamatergic transmission is incorporated to reproduce the transition towards the awake-like asynchronous state, resulting in a linearization of the input-output response function (***Curto et al., 2009***). Alternatively, in our framework the same network arousal leads to a cortical network composed of L5 neuronal assemblies capable of expressing a state-dependent response function (like flip-flop units) which contribute to a further increase in the complexity of the collective dynamics of the awake or awake-like (***D’andola et al., 2017***) brain. Indeed, nonlinear elements together with a topological structure of the network connectivity can further enrich the phase diagram, giving rise at the macroscale to abrupt transitions between dynamical regimes (***Deco et al., 2008; Steyn-Ross et al., 2013***), to hysteresis properties in the speed of the slow waves (***Capone and Mattia, 2017***), and to the enhancement of the sensitivity to the underlying structural organization of the cortical network (***Capone et al., 2019***).

In our case the nonlinearity of the response function is required in order to reproduce the abrupt transition we observed between the two phases in which SO and micro-arousals are alternatively self-sustained. We reproduced this experimental evidence by driving the model to cross these two regions of the bifurcation diagram through a subcritical Hopf bifurcation. In this theoretical framework, our population rate model eventually allowed us to formulate an intriguing hypothesis, namely that micro-arousals and SO in L5 cortical assemblies are two competing attractor states, and as a result the increased persistence in time of one of them is mainly due to the destabilization of the other. According to this hypothesis, we found that the MUA change across the Down-to-Up transition during the SO phase was wider than the activity jump measured during Down-to-micro-arousal transitions (Figure 8A). Indeed, during the latter, the orbit attracting the Up-Down state cycle lost its stability spiraling towards a micro-arousal, with lower firing rate than the Up state (see model trajectories in Figure 7D). For smaller MUA gaps in the Down-to-micro-arousal transition a larger firing rate variability was found (MUA SD, Figure 8C) indicating that the attracting force towards the SO orbit was progressively weakened. The fingerprint of such attractor competition was then a mean-standard deviation anticorrelation of the MUA that we remarkably found in the experiments (compare Figure 7E,F with Figure 8C,D).

According to our *in vivo* recordings, the frequency of the cyclic alternation between SO and micro-arousals we found has a stable fixed value of 0.2 Hz throughout time. Evidence of an alternation between synchronized and desynchronized states under light ketamine-induced anesthesia were previously observed in cat intraparenchymal recordings (***Miyasaka and Domino, 1968; Mori et al., 1971***). Intriguingly a similar phenomenon occurs in humans with ketamine at general anesthesia levels (***Akeju et al., 2016***). In this case, slow-delta oscillations in EEG recordings alternate with periods that display prominent gamma oscillations (“gamma bursts”), similar to what we found during light anesthesia (Figure 1D,E). As the plasma levels of ketamine decrease in both humans and cats, SO dissipate, allowing for prolonged periods of low-amplitude, high-frequency oscillations (“stable beta/gamma oscillations”), matching the elongation of the micro-arousals we observed between L1 and L2. The periodic transition between SO and micro-arousals then appears to be a universal hallmark of the arousal process from ketamine-induced anesthesia in mammals. In rats, we found that this 5 s cyclic pattern of activity is rather regular across time and animals, giving rise to a 0.2 Hz peak in the LFP power spectrum (Figure 1C,D). We expect to see the same spectral feature in the EEG of ketamine-anesthetized humans but at a lower frequency, as gamma bursts have a seemingly slower pace (about a 10 s cycle) (***Akeju et al., 2016***).

In order to reproduce this phenomenon our population rate model of an L5 neuronal assembly had to incorporate an input ***I***_*sin*_ sinusoidally modulated at 0.2 Hz, causing the system to cross back and forth a subcritical Hopf bifurcation. This is a rather general mechanism exploiting the sensitivity to an external perturbation of the network working close to a critical point. We assumed the sinusoidal modulation to be exogenously generated, and as different brain circuits might in principle provide such input during the arousal process, a reasonable expectation is to see the cyclic SO-micro-arousal pattern not only under ketamine-induced anesthesia, but also in other brain states. In partial support of such a hypothesis, a rhythmic alternation between “activated” (low-voltage, fast-activity) periods and “deactivated” SO phases was observed in rats under urethane anesthesia (***Clement et al., 2008; Chan et al., 2015***). The period of these activated-deactivated cycles was relatively long (about 11 min on average) resembling the REM/NREM cycle of natural sleep. Indeed, a stimulation of the cholinergic nuclei in the brainstem, such as the pedunculopontine tegmental nucleus (PPT), which are known to regulate the cyclic brain-state alternations during sleep, systematically disrupted the aforementioned alternating activity pattern (***Clement et al., 2008***). Intriguingly, the causal involvement of the ascending reticular activating system has also been preliminarily proven to interrupt slow-delta oscillations in the cortex of ketamine anesthetized cats eventually giving rise to activated periods (***Miyasaka and Domino, 1968***).

Considering the brainstem as the possible source of such periodic modulation of the cortical excitability which we modeled as a sinusoidal input ***I***_*sin*_, it is tempting to interpret infra-slow oscillations (0.02 – 0.2 Hz) ubiquitously observed during natural sleep (***Vanhatalo et al., 2004***) or resting wakefulness (***Raichle et al., 2001***), as another expression of the cyclic bouncing between metastable SO and micro-arousal attractor states. Indeed, infra-slow oscillations display a spectral power strongly anti-correlated with the magnitude of faster EEG oscillations (> 1 Hz, including gamma band) (***Vanhatalo et al., 2004***), such that a low-***I***_*sin*_ phase would correspond to an enhanced slow-delta power (SO, 0.5 – 4 Hz), whilst a high-***I***_*sin*_ phase would promote an activated state with pronounced high-frequency gamma rhythms.

A remarkable example of infra-slow oscillations is the cyclic alternating pattern (CAP), described as short and repetitive interruptions of the normal sleep oscillatory activity in humans (***Terzano et al., 1985; Parrino et al., 2012***). Within a relatively wide range of possible CAPs, the so-called subtype A1 displays an alternation between SO and activated states with smaller EEG fluctuations reminiscent of what found in ketamine-anesthetized rats. CAP periods range from 20 s to 40 s (***Terzano and Parrino, 2000; Achermann and Borbely, 1997);*** that is, in the 0.025 – 0.05 Hz frequency band, slower than the 0.2 Hz we report here. Notably, CAP has been proposed as a natural marker of sleep instability (***Parrino et al., 2012***), as it often onsets during the transition periods between REM and NREM sleep. Furthermore, administering an arousing stimulus during the activated states of CAP, SO are immediately elicited.

Such susceptibility to external perturbations and the sensitivity to the arousal level are both dynamical features expressed by a cortical network working close to a critical point, as in the one presented here. On the other hand, differences in the pace of the SO-micro-arousal alternation found between natural sleep and anesthesia might be explained by a different mechanistic origin of ***I***_*sin*_. Addressing all these open issues deserves further investigation relying on intracortical activity recordings in natural sleeping animals. Starting from this, we believe that the theoretical and analysis framework we developed here could be an effective tool in identifying the possible common origin of the infra-slow oscillations found during the awakening from natural sleep and deep anesthesia.

## Methods

### *In vivo* LFP recordings

Adult male Wistar rats (271 ±31 g, *n* = 5) were anesthetized via an intraperitoneal injection of ketamine (60 mg/kg) and medetomidine (0.5 mg/kg) (***Bettinardi et al., 2015***). Atropine (0.05 mg/kg) was injected subcutaneously to prevent secretions. All pressure points and tissues to be incised were infiltrated with lidocaine before surgery. Body temperature was maintained at 37°C using a water-circulating heating pump. Electroencephalogram and electrocardiogram were monitored during the experiment. Heart rate values remained stable between 290 and 300 bpm.

In order to compare experiments that had different durations we selected four intervals of 10 min at different levels of decreasing anesthesia (see Figure 1A). In each experiment, we took Deep1 (D1) as the first 10 min of recording, starting 44 ± 2 min after induction on average. The appearance of the first micro-arousal occurred, on average, 185 ± 35 min after induction. Deep2 (D2) was identified as the last 10 min of pure SO before that first micro-arousal, and the following 10 min corresponded to the interval Light1 (L1). As soon as waking signs appeared (withdrawal reflex, elevated heart rate, fast breathing) a reinduction dose was administered to return to the initial deep level of anesthesia. The last 10 min of recording before reinduction time were identified as Light2 (L2).

LFP recordings (by LFP here we refer to the unfiltered signal) were obtained from the primary visual cortex, V1 (−7.3 mm antero-posterior, 3.5 mm medio-lateral) of the right hemisphere. All coordinates are relative to Bregma (***Paxinos and Watson, 2004***). A 16-channel silicon probe (1 shank with 16 linearly spaced sites at 100 *μ*m increments with impedances of 0.6–1 ***M*Ω** at 1 kHz, from NeuroNexus Technologies) was introduced perpendicular to the V1 cortex under visual guidance, until the most superficial recording site was aligned with the cortex surface (***Mattia et al., Under review***). LFP recordings (in mV)were amplified with a multichannel system (Multi Channel Systems). The signal was digitized with a CED acquisition board and Spike2 software (Cambridge Electronic Design).

All experiments were supervised and approved by the University committee (Universitat de Barcelona) and complied with the European Convention for the protection of vertebrate animals used for experimental and other scientific purposes (Strasbourg 3/18/1986) and the local law of animal care established by the Generalitat de Catalunya (Decret 214/97, 20 July).

### Data analysis

LFPs from different layers of V1 were simultaneously acquired at a sampling frequency of 9921 Hz during the fading of anesthesia and saved in consecutive 900-s-long recordings.

MUA was estimated as the power of the Fourier components at high frequencies (200 – 1500 Hz) of the extracellular recordings (LFP) in windows of 50 ms (***Reig et al., 2010; Sanchez-Vives et al., 2010; Ruiz-Mejias et al., 2011***). MUAs were logarithmically scaled and their values were shifted in order to center the peak corresponding to the Down state at *log*(***MU A***) = 0. As shown in ***Mattia and Del Giudice (2002)***, this time series provides an accurate estimate of the population firing rate because normalized Fourier components at high frequencies have densities proportional to the spiking activity of the involved neurons.

Up and Down states were singled out by setting a minimum state duration of 80 ms and a threshold in log(MUA) values at one-third of the interval between the peaks of the bimodal distribution of log(MUA) corresponding to the Up and Down states. Asymptotic firing rate during Up states was computed as the mean log(MUA) during the Up state, avoiding the first and last 50 ms, corresponding to the transitions from and towards the Down states. States shorter than 150 ms were discarded from asymptotic firing rate computation. For each experiment, we selected the channel with maximum asymptotic firing rate during the Up state, whose location in depth corresponds to cortical layer 5 (***Mattia et al., Under review***).

The moving variance of the LFP (mvLFP) was computed as the variance of the LFP in windows of 1 s with 0.1-s steps in the channel with the maximum asymptotic firing rate. For each experiment, a threshold was set at 15% of the distribution of the mvLFP values of those recordings with pure slow oscillatory activity. In each 1-min time window, LFP data points below this threshold were classified as micro-arousals. Power spectra were computed using the Welch method, and spectrograms were computed in non-overlapping windows of 1-min using the LFP signal high passed at 0.01 Hz.

The evolution across the decreasing levels of anesthesia of duration and firing rate (MUA) of the different states (Up, Down, micro-arousal) was evaluated by averaging the values of state duration or firing rate in time windows of 1 minute across the time of each experiment. To compare between subjects, we used the average value of duration or firing rate of the states that fell into the time intervals selected as D1, D2, L1 or L2 in each experiment.

Comparisons between different levels of anesthesia and statistical significance was set at *p* < 0.05. All analyses were performed using MATLAB (The MathWorks).

### MUA normalization and average over experiments

In order to understand the dynamical nature of micro-arousals, we evaluated the average profile of the log(MUA) around the Down-to-micro-arousal transition over all micro-arousals in each recording. In order to average over different recordings, we rescaled the log(**MUA**) to have its minimum as 0 and its maximum as 1

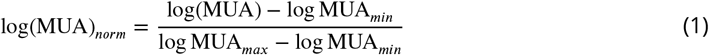

where log **MUA**_*max*_ and log **MUA**_*min*_ are respectively the maximum and the minimum of the profile averaged over all MA states in a single experiment. Once normalized, log(***MUA***)_*norm*_ was averaged over all recordings (Figure 7A in orange). In each experiment, micro-arousals were sorted by level of activity in the Down state before the micro-arousal onset and grouped into three clusters of the same size. In each cluster the average profile of the normalized log(MUA) was computed with Eq.1 and using the same log **MUA**_*max*_ and log **MUA**_*min*_ as estimated above. Finally, the average profile around the Down-to-Up transition from the L1 to L2 periods was estimated (Figure 7A in black) and normalized.

As an indicator of the attractor state stability we resorted to the inter-state variability of the normalized log(MUA) as follows:

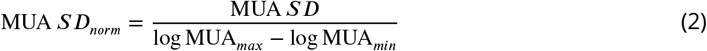

In this analysis, one out of five experiments was rejected as the MUA signal was extremely noisy, probably due to the electrode placement. This led to profiles of single micro-arousals that were not good enough for a dynamical fine structure analysis.

### Population rate model

We considered the mean field dynamics of a population of N excitatory neurons strongly (recurrently) connected. As shown in Figure 5A, the network dynamics also take into account an activity-dependent self-inhibition (adaptation) ***a***(*t*) whose weight is defined by the coupling variable ***g***_*a*_ and an external input current, ***I***_*ext*_, modeling the average excitatory synaptic input provided by presynaptic neurons external from the modeled network. We also considered an additional oscillating input current that, under suitable dynamical conditions, can be turned on to reproduce the alternation between micro-arousals and SO observed under light anesthesia. This oscillatory external input is defined as

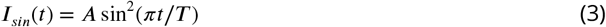

where ***A*** = 1.8 and ***T*** = 2 s are the amplitude and the period of oscillation chosen to fit the experimental observations. The dynamics of the population can be defined by the following coupled equations:

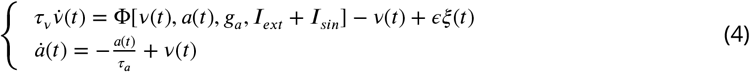

where Φ is the input-output gain function for the *Leaky Integrate-and-Fire* (LIF) neuron (***Ricciardi and Sacerdote, 1979***), ***a***(*t*) is the adaptation variable, ***v***(*t*) is the firing rate, ***ξ***(*t*) is a Gaussian variable describing a white noise of size e. ***v***(*t*) was modeled as a stochastic variable since finite-size fluctuations were also incorporated. Decay times for ***v*** and ***a*** are respectively ***τ***_*v*_ = 20 ms and ***τ***_*a*_ = 500 ms.

The dynamics can be modulated by changing the strength ***g***_*a*_ of the adaptation, and the external current ***I***_*ext*_. Indeed depending on the parameters value the system can be characterized by different configurations of stable fixed points or limit cycles (see Results section). To achieve such analysis standard tools for dynamical models were used (***Strogatz, 2014***). If we consider ***I***_*sin*_(*t*) = 0, depending on the values of ***g***_*a*_ and ***I***_*ext*_, the dynamics of the system can be set in an oscillatory (SO) regime or in a stationary regime with one fixed point, with low or high levels of activity: LAS or HAS regions, respectively (Figure 5B). Finally, the ***I***_*sin*_(*t*) is required in order to have the regular alternation between SO and HAS dynamics observed in experiments.

## Acknowledgments

This work was financially supported by the Spanish Ministry of science BFU2017-85048-R and by CERCA Programme / Generalitat de Catalunya, and by the EU Horizon 2020 Research and Innovation Program under HBP SGA1 (grant no. 720270) and HBP SGA2 (grant no. 785907) to MVSV and MM. We thank C. Gonzalez-Liencres and T. Donegan for editing assistance and P. del Giudice and M. Massimini for useful discussion.

